# Involuntary breathing movement pattern recognition and classification via force based sensors

**DOI:** 10.1101/2022.07.12.499777

**Authors:** Rajat Emanuel Singh, Jordan M. Fleury, Sonu Gupta, Nate P. Bachman, Brent Alumbaugh, Gannon White

**Affiliations:** Department of Kinesiology, Northwestern College, Orange City, IA, USA; Department of Kinesiology, Colorado Mesa University, Grand Junction, CO, USA; Department of Computer Science, Northwestern College, Orange City, IA, USA

**Keywords:** Acceleration, Jerk, Involuntary Breathing Movement, Pattern Recognition, Classification

## Abstract

The study presents a novel scheme that recognizes and classifies different sub-phases within the involuntary breathing movement (IBM) phase during breath-holding (BH). We collected force data from eight recreational divers until the conventional breakpoint (CB). They were in a supine position on force plates. We segmented their data into the no-movement (NM) phase aka easy phase and IBM phase (comprising several events or sub-phases of IBM). The acceleration and jerk were estimated from the data to quantify the IBMs, and phase portraits were developed to select and extract specific features. The K means clustering was performed on these features to recognize different sub-phases within the IBM phase. We found five-six optimal clusters separating different sub-phases within the IBM phase. These clusters separating different sub-phases have physiological relevance to internal struggle and were labeled as classes for classification using support vector machine (SVM), naive bayes (NB), decision tree (DT), and K-nearest neighbor (K-NN). In comparison with no feature selection and extraction, we found that our phase portrait method of feature selection and extraction had a low computational cost and high robustness of 96–99% accuracy.

## 1. Introduction

There are involuntary physiological responses present in the biological systems that are activated when life-threatening situations arise. The prolonged contraction of the diaphragm and external intercostal muscle during BH results in one such response. It is considered that prolonged contraction and hypercapnia during BH activate chemoreceptors [1] [2]. These receptors reaching the sensory cortex generate a dyspnea signal, which is crucial for the initiation of breathing [3] [4]. But, humans extend BH through the phasic involuntary diaphragmic contractions, especially under certain circumstances such as free diving [5].

The question is whether sensors can detect movement patterns associated with these involuntary contractions? The question is crucial for abnormal breathing detection, and the answer is in the duration and strength of involuntary muscle contractions during the struggle phase (SP). SP is a phase in which IBMs appear due to deep muscle involuntary contractions. Thus, invasive methods such as pressure transducer-based catheters can capture them [6], but non-invasive methods such as surface EMG, and force plates during strong contractions can be applicable [4], [7], [8]. The IBM starts with the physiological breakpoint ((PB), the point where the first IBM appears) and ends with CB (the point where breathing is resumed voluntarily) [7]. The IBMs are periodic and very subtle and are controlled autonomously during SP [4]. Thus, rhythmic varying patterns of IBMs associated with involuntary periodic contractions can be acquired using force based non-invasive methods.

Previous studies used a simple computational model to detect phases during BH (onset and end offset of the IBM phase) [6]. However, the IBM phase is more complex than just onset and end offset. IBMs in SP are rapid continuous movements of varying magnitude. Previous studies had shown complex varying phases of IBMs, as IBMs’ amplitude and frequencies kept increasing till the CB [9], [10]. Thus, using a simple model, it is hard to detect patterns or sub-phases in a movement of such nature. Moreover, the current literature does not present methods to recognize such sub-phases within the IBM phase. Therefore, methods emphasizing detecting patterns or sub-phases within the IBM phase are of importance in clinical science and should be developed. Hence, the goal of our study is to design such a method. We refer to the IBM phase as several events or patterns of IBMs during the SP.

In this study, we present a novel scheme (shown in figure 1) that can recognize different sub-phases within the IBM phase during BH, and classify them accordingly. The force sensors were used to capture kinetics associated with IBMs as it comprises mechanical components [4]. We did not use force data directly for the categorization because kinematic patterns of rapid movements can be quantified nicely through jerk and acceleration. Hence, the rapid movements related to involuntary contractions during BH was estimated using acceleration and jerk. These outputs were later used as features for pattern recognition and classification of IBM sub-phases.

**Figure 1.**
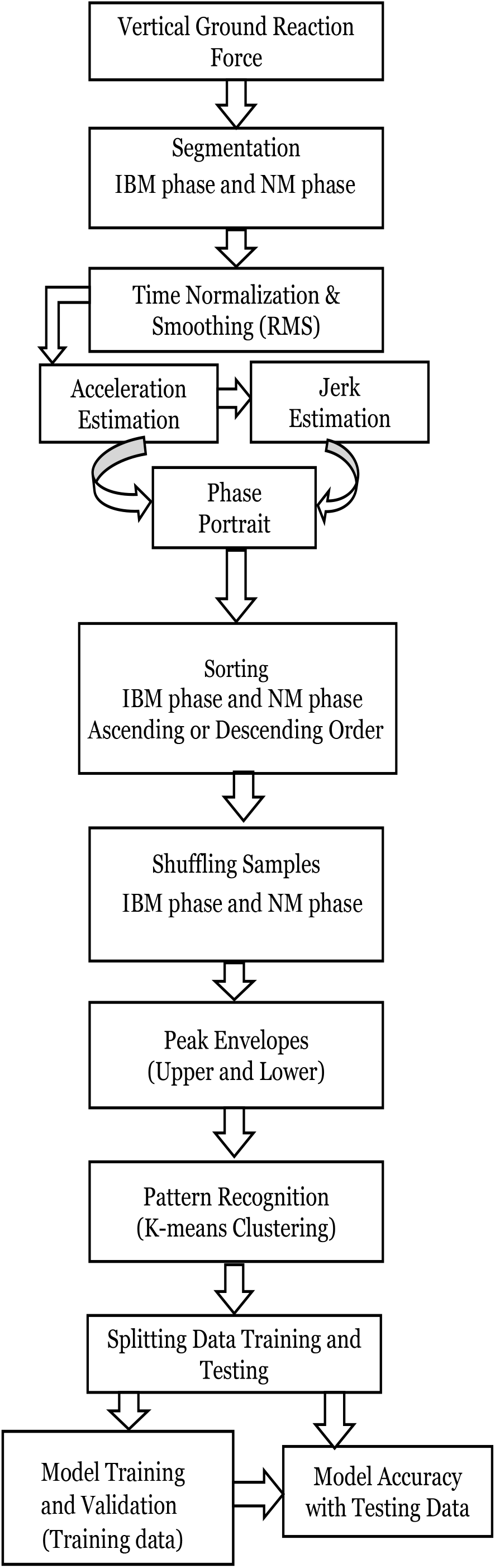
The framework for the clustering and classification of IBM phases.

## 2. Materials and Methods

The IRB review board of Colorado Mesa University has approved this study. In this study, we recruited eight healthy participants who are recreational divers. Divers generally have the higher lung capacity and better control of their breathing due to years of intense training. This makes them a better fit for our study. The demographic data of participants recruited in our study is listed in table 1.

**Table 1.** Demographic data of participants

| *Participant ID* | *Sex* | *Age<sub>u</sub>* | *Height<sub>v</sub>* | *Mass<sub>w</sub>* | *Experience<sub>u</sub>* |
| --- | --- | --- | --- | --- | --- |
| P1 | Male | 20s | 183.5 | 87.5 | 6 |
| P2 | Female | 20s | 174.5 | 72.0 | 6 |
| P3 | Male | 20s | 183.0 | 78.0 | 15 |
| P4 | Male | 20s | 171.0 | 67.0 | 1 |
| P5 | Female | 20s | 164.0 | 63.5 | 12 |
| P6 | Male | 20s | 190.0 | 79.0 | 2 |
| P7 | Male | 20s | 174.5 | 89.5 | 3 |
| P8 | Male | 20s | 189.0 | 114.0 | 5 |

Units:- *u* years, *v* centimeters, *w* kilograms, *u* years.

### 2.1. Experimental Design

The protocol to collect data from a single participant required a single visit to our lab. On the visit, signed consent was obtained, and data regarding their age, gender, height, weight, and years of experience was recorded as shown in table 1. They were also instructed to avoid eating, drinking, and exercising two hours before the data collection. Two AMTI force plates were used to acquire the data from the participants at a sampling rate of 1Khz. We asked the participant to lay in a supine position on the force plates. The participants were laying with a static posture and covered the force plates surface mostly with their upper and middle back. Moreover, to maintain a comfortable posture during data collection, a cushion and a rolled-up pad were positioned under their head and knee.

We introduced a warm-up session in three steps. In the first step, participants breathed normally (relaxation) for five minutes followed by a minute of BH. In the second step, the relaxation period was reduced to two minutes, and the BH phase was incremented by an additional sixty seconds from previous step. In the third step, participants relaxed again, followed by three minutes of BH.

Prior to data collection participants were given enough relaxation periods. For data collection, they were asked to hold their breath as long as they can. The data was recorded from the participant’s first inhalation (starting BH) to voluntary exhalation (ending BH). The participants were asked to lay down after data collection until lightheadedness subsides.

### 2.2. Data analysis

We used R programming software (R 4.2.0) for signal processing and data analysis. The supine posture and the muscle spasm exert force vertically resulting in the appearance of IBMs primarily in the vertical direction (z). Hence, force plates were used to capture this biological phenomenon, and the force data in the vertical direction (vertical ground reaction force) was analyzed.

The IBMs appeared later during the BH. Thus, we segmented vertical ground reaction force data into the NM phase and IBM phase through visual inspection. The NM phase data comprised mostly of baseline noise. The IBM phase comprise of several events of IBMs. Hence, it was easy to isolate these two phases with visual inspection.

The segmented data was time normalized (spline interpolation) to 10000 time points for the IBM phase (10000 × 1) and NM phase (10000 × 1) for each participant (10000 × 1 × 8). Later the segmented data was smoothed using a root mean square function with a window size of 50 ms. The IBM and NM data (time normalized and smoothed) were combined to form a processed vertical ground reaction force data vector (20000 × 1 × 8).

### 2.3. Acceleration and Jerk estimation

The participants had no significant center of mass (*COM*) displacement of other body segments due to their static posture. Therefore, we assumed full body *COM* in the chest, and acceleration 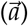 and jerk 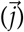 of the IBM phase were estimated from the processed vertical ground reaction force (*F*_*z*_), as shown below.

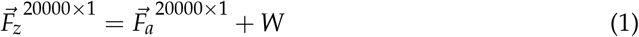

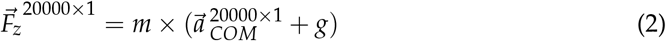

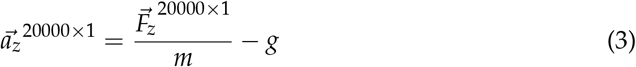

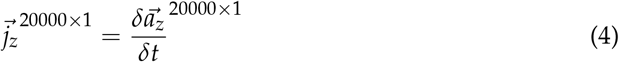

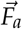 is the dynamic force data vector due to the accelerating center of mass 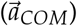, *W* is the weight of the participants’ chest, *m* is the mass of the participant’s chest. The participants’ chest mass is calculated from the thorax percentage contribution to full-body mass [11], and *g* is the acceleration due to gravity. The superscript displays the dimension of the data vector for a single participant in equations (1) to (4).

### 2.4. Phase portraits

A phase portrait 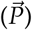 is a geometric representation of the dynamic trajectories in the phase plane [12]. The 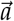 and 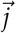 data were used to develop phase portraits for feature selection and extraction. The phase portraits yielded phases that overlap between the IBM phase and the NM phase. Thus, developing phase portraits were crucial to remove phases and frequencies, that overlap between the NM phase and IBM phase. This feature selection and extraction through phase portraits may pose useful to enhance the performance (accuracy, computation, robustness) of our pattern recognition and detection scheme.

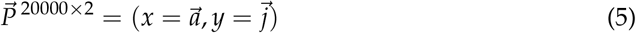

We first separated the data points overlapping between IBM phase and NM phases by sorting the values of 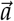and 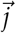 in descending/ascending order from phase portraits. These samples of 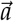 and 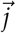 were then shuffled randomly and independently so that the features will represent unbiased population of the data. Furthermore, peak envelopes, lower 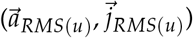 and upper envelopes 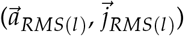 were extracted. These features were finally used for unsupervised and supervised learning.

### 2.5. Unsupervised learning

We performed unsupervised learning to recognize similarities and/or dissimilarities between the IBM and NM phases, and to recognize specific patterns/sub-phases/groups within IBM phase. These groups were labeled as different classes for classification.

K means cluster analysis was implemented on the extracted features 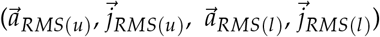 as an unsupervised learning algorithm. The K means clustering algorithm groups the data points within these features based on their similarity into clusters.

### 2.6. Statistical analysis

To test the normality, we performed Kolmogorov–Smirnov test. The type (parametric or non-parametric) of test to compare the difference between the IBM phase and NM phase was based on the rejection of the null hypothesis. We considered a significance value of *p* < 0.05 to reject the null hypothesis.

Violin and box plots are used to display the distribution, mean and standard deviation of the data.

### 2.7. Supervised learning

A supervised learning approach was implemented to isolate the onset of different sub-phases of IBMs with a classifier. We developed a classification model with training data that makes no assumption about the normality of data. We used SVM, NB, DT, and K-NN as classifiers.

The features that were grouped into different clusters (NM phase and different IBM sub-phases) were labeled as different classes for supervised learning. Furthermore, these labeled features were split into a ratio of 75:25 for training 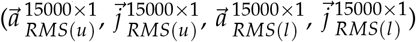 and testing 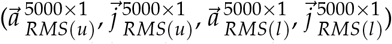 data. The training data was used to train the classification model, and the test data was used to evaluate the accuracy of the model. We also used K-fold cross (K = 10) validation procedure to validate our training model before evaluating its accuracy with testing data.

## 3. Results

## 3.1. IBM phase vs NM phase

We first tested whether the IBMs were present in the vertical ground reaction force, and were detectable through test statistics. We found vertical ground reaction force data was statistically significant (*p* ≤ 0.05, Wilcoxon sign rank paired test) between the IBM and NM phases. The force amplitude’s median value were different between the IBM phase and NM phase as shown in figure 2.

**Figure 2.**
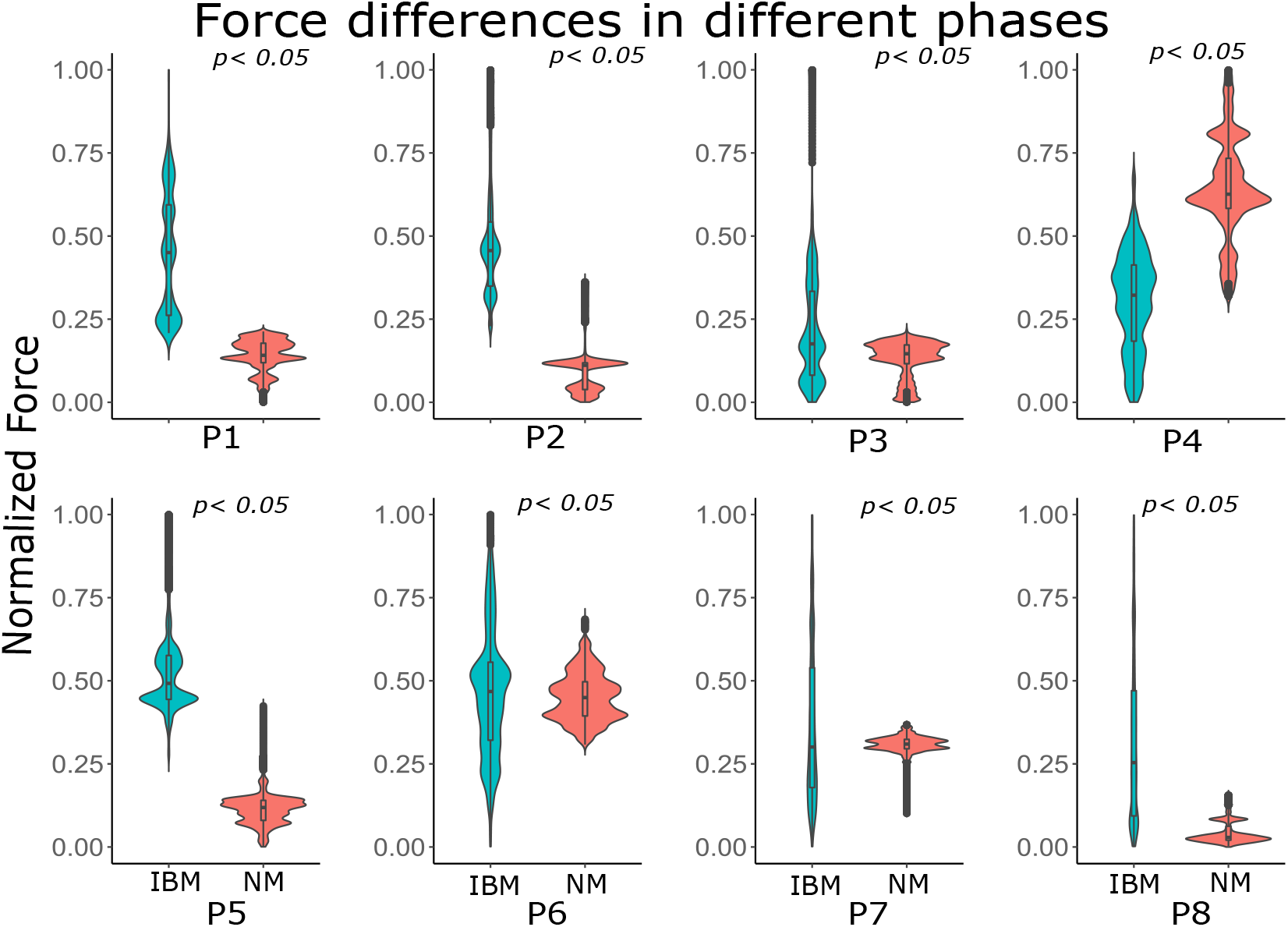
The top panels labeled with participants’ ID (P1, P2, P3, P4) and the bottom panels labeled with participants’ ID (P5, P6, P7, P8) show the violin plots of the respective participants. The y-axis is normalized force value. A box plot is also encapsulated within the violin plot. The grey region represents the high spikes (outliers) in the data, and the *p* values for each participant has been displayed on the top of each panel.

Moreover, statistically significant differences (*p* ≤ 0.05, Levene’s test) in the variance of the ground reaction force between the NM phase and IBM phase were present for all the participants. Figure 2 shows the distribution of the vertical ground reaction force data for all the participants.

The high variance during the IBM phase was due to periodic signal output of varying amplitude. Hence, the IBMs of varying magnitude were present in the force data and were detected using an appropriate statistical measure.

## 3.2. Phase portraits

The phase portraits generated from the estimated 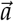 and 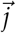 were then used for feature selection and extraction. We found clear isolation between the IBM phase and NM phase from the phase portraits and their density plot, as shown in figure 3.

**Figure 3.**
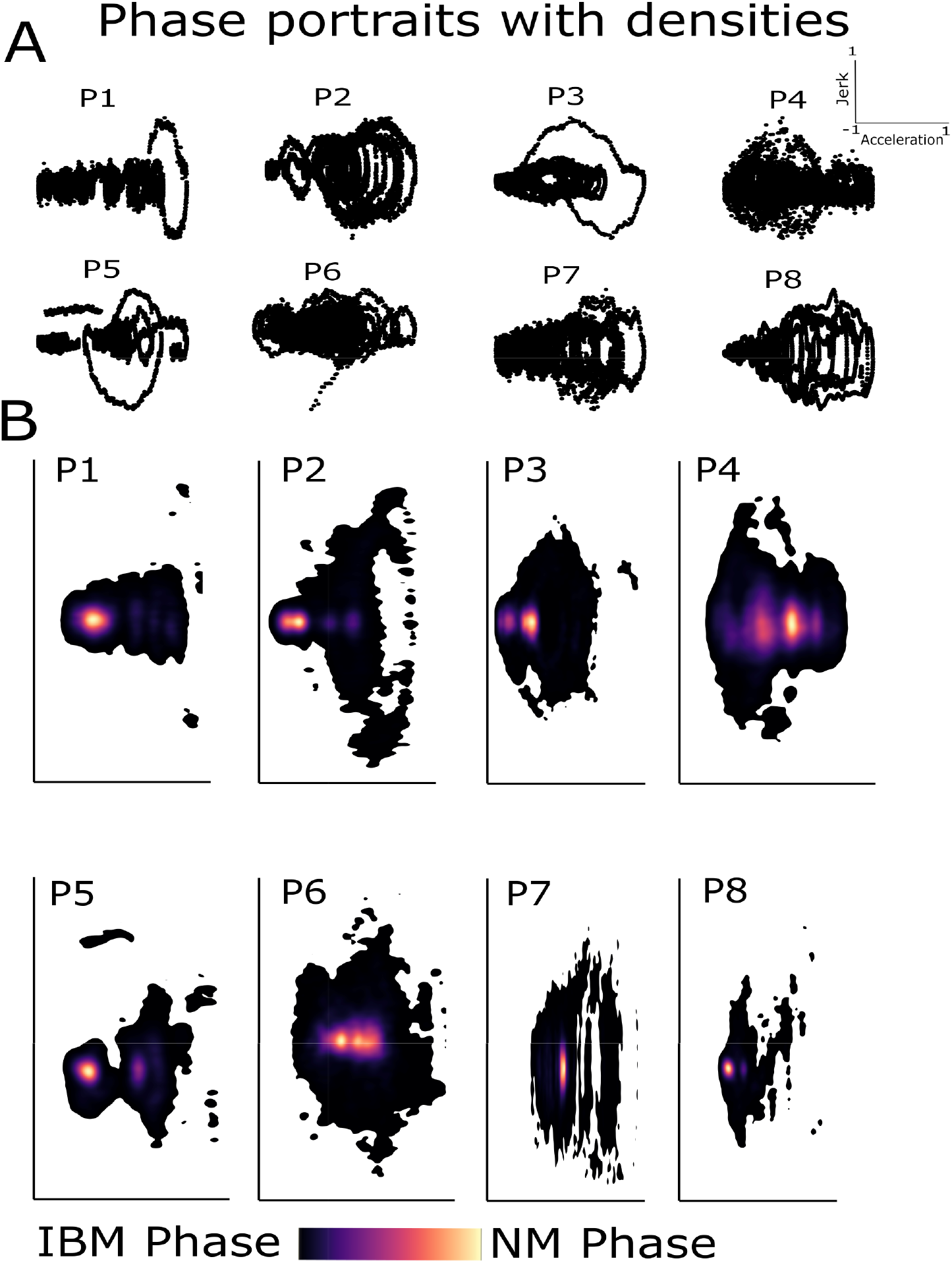
A) The top eight panels display phase portraits for all the participants. Thus, each panel shows a phase portrait of a participant. The phase portrait indicates rhythmicity of varying magnitude (amplitude) during IBM phase. B) The bottom eight panels show the density of data points in the phase portraits. Thus, each panel displays the density plot of a participant. The NM phase and IBM phase isolation are visible. The 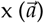 and 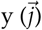 axes values are normalized between (−1,1) across participants for better visual representation. A spectrum of densities (bottom) can be observed during the IBM phase which is associated with different sub-phases during SP.

The data points for the NM phase were concentrated at a specific region of less magnitude around the baseline signal. The data density was also high in this region as shown in figure 3. Thus, the BH data was mostly composed of baseline noise associated with the NM phase.

The IBM phase concentrically surrounds the NM phase area. These concentric circles represent the regions of low, moderate, and high magnitude of 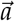 and 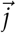 as shown in figure These different regions within the IBM phase suggest different sub-phases or patterns within the IBM phase. In addition, the circular shape of the phase portrait suggests the presence of some rhythmic patterns of IBMs.

### 3.3. Clustering

The goal of clustering is to identify sub-phases within the IBM phase. But before, we first determined the optimal number of clusters (K) that need to be defined in the K means algorithm. We found that based on total withinness, five to six clusters were an optimal choice for the goodness of clustering across most participants.

We then tested through clustering whether the IBM phase is a biological phenomenon comprising varying magnitude of 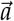 and 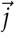 We extracted peak minima and maxima values of the NM phase and IBM phase (sliding window = 25 samples) from phase portraits before feeding them to K means algorithm. The cluster analysis displays clear isolation between the NM phase and IBM phase. Moreover, it also shows the separation of sub-phases within the IBM phase.

The cluster analysis displayed that cluster 1 and cluster 6 comprise NM phase and few IBM data points. The centroid of these clusters was towards the point of origin (0,0) and 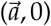. Therefore, suggesting an overlap between the NM phase and IBM phase due to no activity. The values slight above (0,0) for 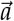 and 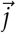 in cluster 1 and 6 suggest IBM onset. Moreover, the constant 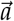 of higher magnitude was also grouped in clusters 1 and 6.

The cluster 2, 3, 4 and 5 mainly represented IBM phase. The centroid of the clusters 2, 3 4 and 5 showed varying degree of 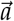 and 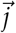 during IBM phase as shown in figure 4. These clusters explained phases of 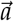 and 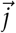 with (high, high), (high, moderate), (moderate, high) and (moderate, moderate) magnitude. The categorical names here were based on the relative magnitude of the centroids. Thus, different patterns or sub-phases were present and identified during IBM phase using K means clustering.

**Figure 4.**
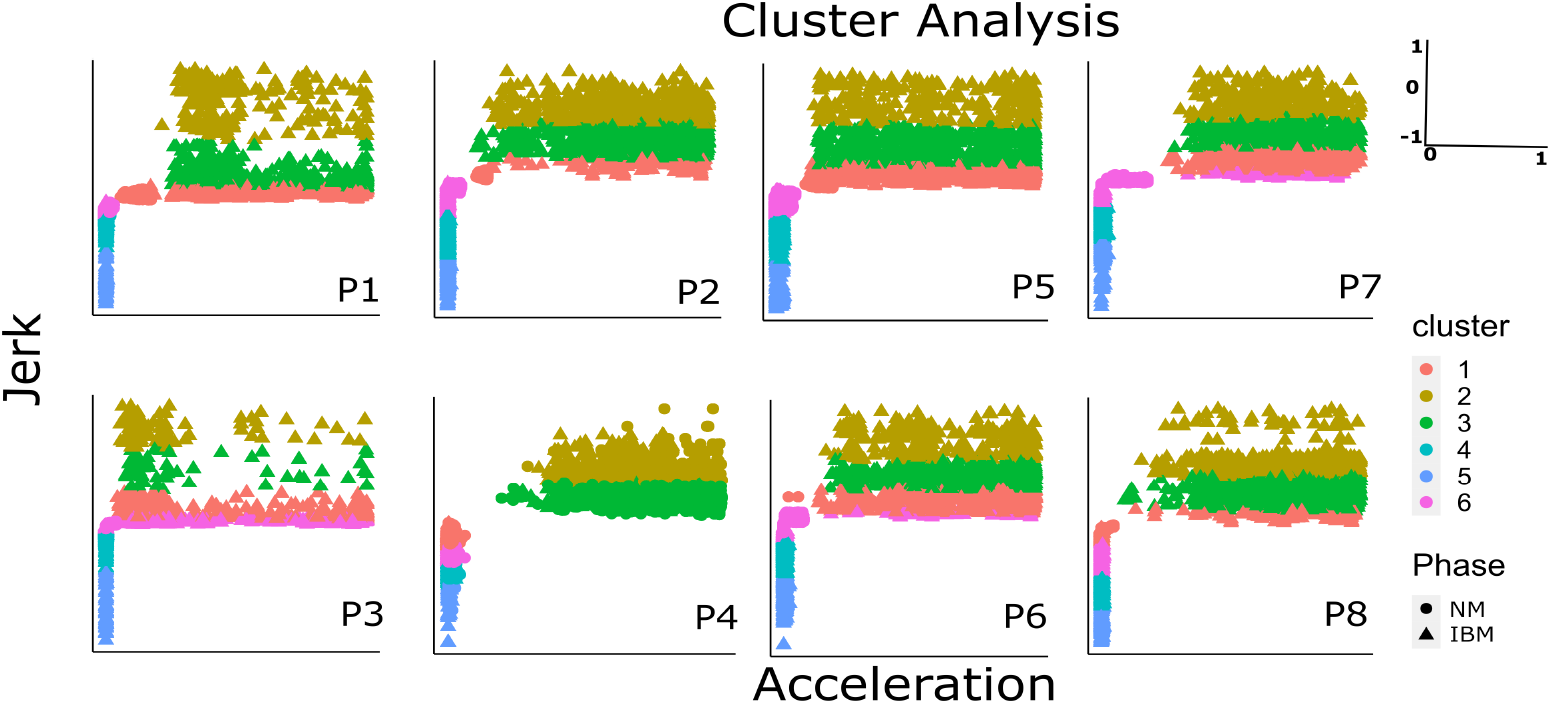
K means cluster analysis for all the participants. Each panel shows the results of K means clustering for a single participant. The magnitude for 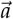 on the x-axis and 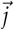 on the y-axis was normalized between 0 to 1, and −1 to 1 respectively. The NM phase and IBM phase is represented by circle and triangle shape, respectively. Cluster 1 and 6 shows overlap between NM and a certain IBM sub-phases, whereas cluster 2, 3, 4, and 5 shows more patterns revealing different sub-phases of IBM.

### 3.4. Classification

Supervised learning classified these complex patterns or sub-phases with a computational model. Our aim of classification was to design a scheme that detects the clustered regions with high accuracy.

We trained different classifiers for the labeled data. The data were labeled based on the clusters. We performed 10-fold cross-validation procedure to validate our classification model (shown in figure 5A). We further evaluated the model accuracy with the testing data. We found that our classification model can predict with an accuracy of 96.5-99.9%.

**Figure 5.**
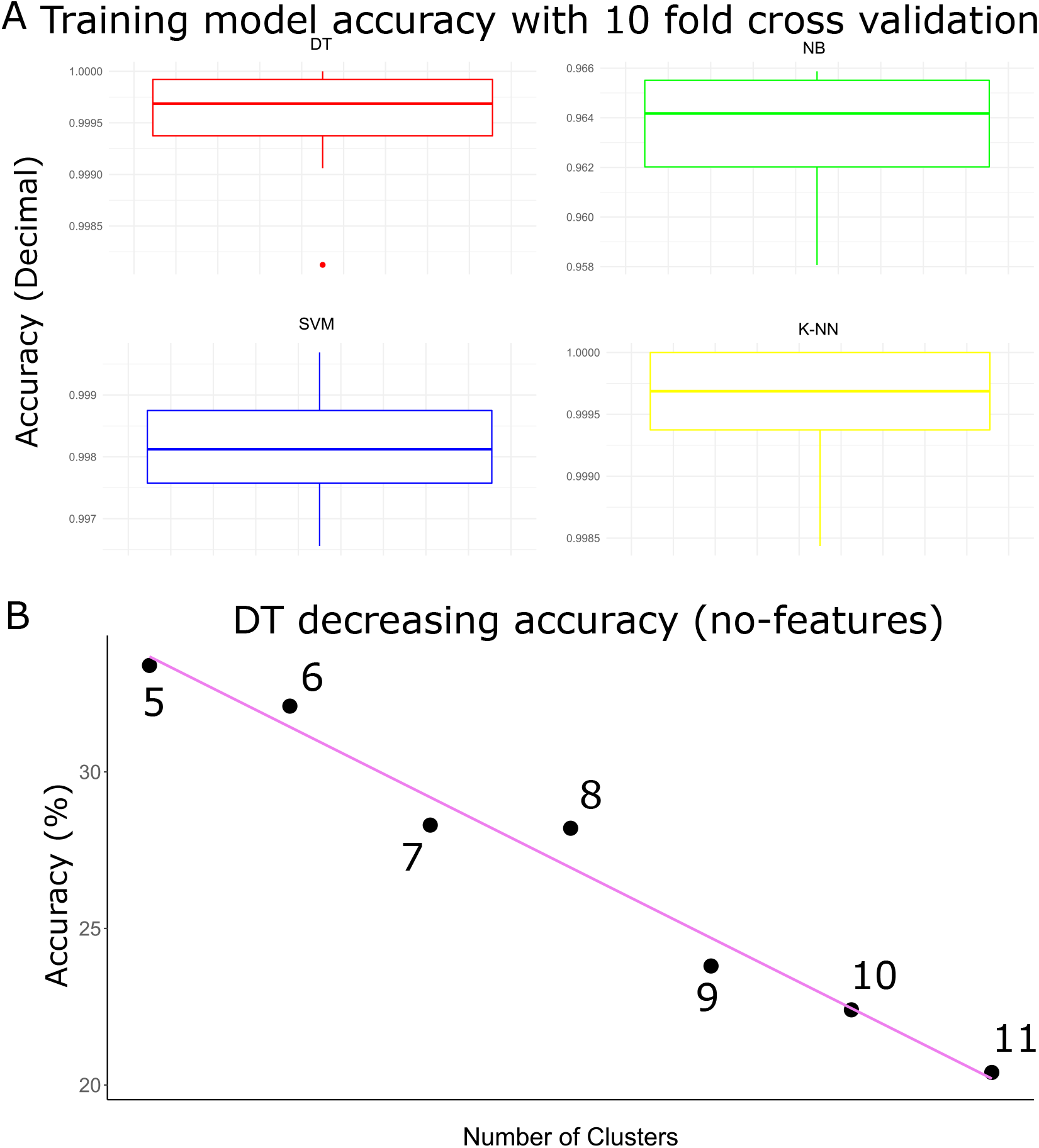
A) The top four panels with boxplots show classifiers’ performance using 10-fold cross-validation, thus, each panel displays a specific classifier. The y-axis shows validation accuracy in decimal. The features were selected and extracted to train the model from 5-11 classes, and a 10-fold cross-validation was performed before performance evaluation with testing data. The classes number were also changed to test the sensitivity of accuracy. B) The lower panel shows the relationship between the number of classes and the accuracy of the DT classifier on data without features. The x-axis shows a number of clusters and the y-axis shows accuracy in percentage. The DT accuracy decreased when the number of clusters or classes increased, especially when features were not selected and extracted.

We also trained models without feature selection and extraction. We found that the accuracy of those models was consistent with most algorithms, except DT as shown in table 2. The DT accuracy was also sensitive to the number of classes when no features were selected and extracted as shown in figure 5B. The processing speed also slowed down for most algorithms due to the increased computation cost, as shown in table 2.

**Table 2.**
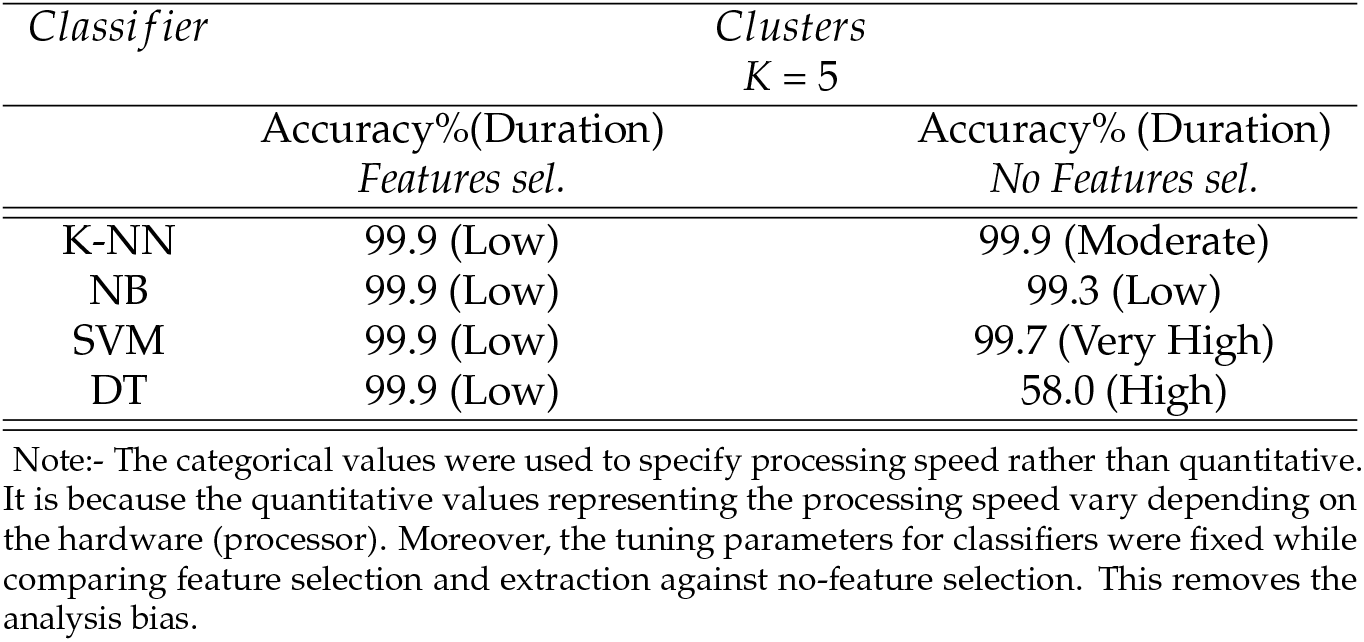
Comparison of accuracy of different methods evaluated using testing data.

| *Classifier* | *Clusters*<br>*K = 5* |  |
| --- | --- | --- |
|  | Accuracy%(Duration)<br>*Features sel.* | Accuracy% (Duration)<br>*No Features sel.* |
| K-NN | 99.9 (Low) | 99.9 (Moderate) |
| NB | 99.9 (Low) | 99.3 (Low) |
| SVM | 99.9 (Low) | 99.7 (Very High) |
| DT | 99.9 (Low) | 58.0 (High) |

Note:- The categorical values were used to specify processing speed rather than quantitative. It is because the quantitative values representing the processing speed vary depending on the hardware (processor). Moreover, the tuning parameters for classifiers were fixed while comparing feature selection and extraction against no-feature selection. This removes the analysis bias.

However, the feature selection reduced the computation cost without loss of accuracy (robustness). In addition, we also found feature selection improves DT accuracy, as shown in table 2. Therefore, our scheme provided a low computation cost with feature selection and higher accuracy with different classifiers. Thus, our model can classify or detect different sub-phases during IBM phase with high accuracy using different algorithms, and low computation cost, as shown in table 2.

## 4. Discussion

We developed a pattern recognition and classification scheme to detect different sub-phases of IBMs during BH. We used force plates and estimated 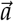 and 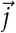 to quantify the movement during contractions. Furthermore, we used phase portraits as a means to select features for our clustering and classification algorithms. We found that our designed scheme has high processing speed, robustness, and accuracy using classifiers such as SVM, NB, DT, and K-NN. We suggest that our scheme is of significant interest to practitioners working with the breathing disorder population. It can assist in detecting certain events of varying intensities within the IBM phase, thus diagnosing the extent of the breathing problem.

The use of kinetic, kinematics and neural signals (EMG) is not uncommon for movement classification [13], [14], [15]. Based on the application of our study, the duration and strength of IBMs, kinetics, and kinematics are better choices for signal classification. A sensor review study validates it [17]. Moreover, surface EMG (sEMG) based classification was challenging because the muscles associated with IBMs are mostly deep (intercostals and diaphragm).

The circular geometry estimated from 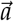 and 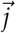 phases during BH suggest rhythmic kinematic patterns of IBMs. There are phase portrait-based comparative studies performed on the locomotion of patients with Parkinson’s and without Parkinson’s that validate this hypothesis [18]. Although previous studies had focused on movements where joint angle during muscle contraction changed [19], our study is the first to use phase portraits for involuntary contraction analysis. Moreover, the variability in the IBMs amplitude appeared as the concentric circles in the phase portraits. Thus, the phase portraits provided a good indicator of periodicity in signals with varying amplitude. And phase portraits provided information about redundant features and crucial features. The redundant features were removed, whereas crucial features were extracted for cluster analysis. The processing, selection, and extraction of features using phase portrait increased the classification accuracy and reduced the computation burden.

### 4.1. Physiological interpretation of clusters

We used clusters to group different phases of IBM phase. These groups displayed changes in 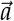 and 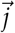 data values during a BH. We found from these clusters that there are sub-phases of low, moderate, and high magnitude within the IBM phase. The low values of 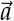 and 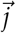 during IBMs were clustered with NM phase in clusters 1 and 6. The low values of 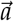 and 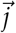 in these clusters represented the onset of IBM. Moreover, the high and moderate data values of IBM phase grouped in clusters 2, 3, 4 and 5 has physiological relevance to extend BH.

Our study showed different levels of 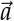 and 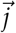clustered in groups 2, 3, 4, and 5 were correlated with force and were associated with increasing IBMs or struggle phase during BH. The struggle phase is a phase in which participants can hold their breath beyond PB before reaching CB [10]. A previous study showed an increased amplitude and frequency of pressure signals is an indicator of struggle termination [6], [7], [10]. In our study, we observed similar results for most participants, where force amplitude incremented continuously until the end of the SP, as shown in figure 6B. However, in a few participants, the signal amplitude decayed before the SP ended, as shown in figure 6A. The low force signal amplitude before SP end could be due to high fatigue index and low energy expenditure as the *O*_2_ level in blood during BH was depleting [20], [21].

**Figure 6.**
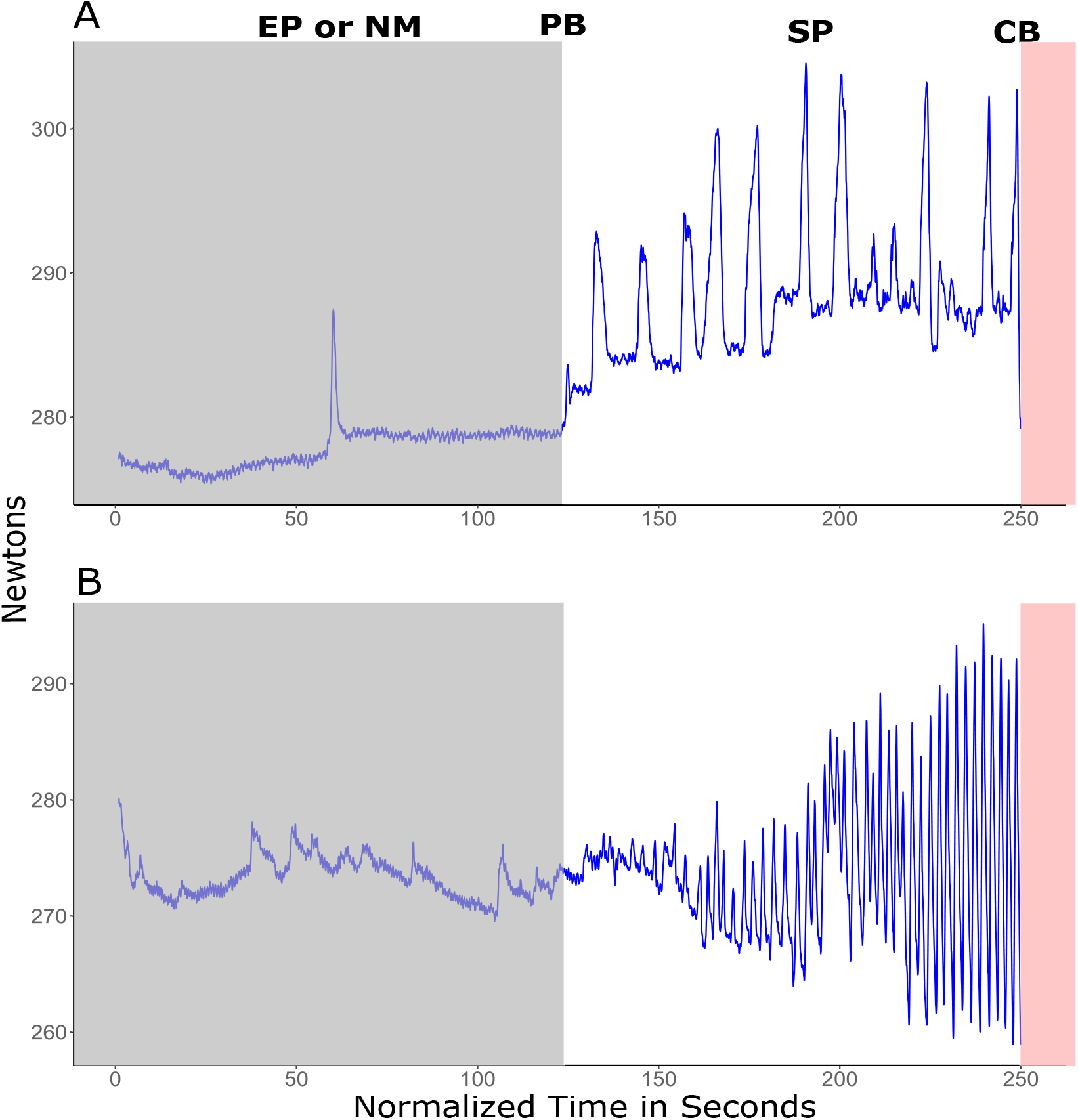
Raw force signal during BH. The top panel (A) and bottom panel (B) shows force signals for two different participants. The x-axis is normalized time in seconds. The time was normalized across participants for better visual representation. The y-axis is force magnitude in newtons which is not normalized here. The grey region shows NM or EP phase, the white region shows SP, and the red region starting from the CB shows end of BH. The spikes appearing during BH is a movement artifact due to some position change on force plate which were later removed from the analysis. A) The force signal for a participant displaying slight decrease in amplitude near the end of SP. B) The force signal for a participant showing continuously increasing amplitude until the end of SP.

### 4.2. Accuracy, Speed, and Robustness

We used different classifiers that makes no assumption about normality, such as SVM, DT, NB, and K-NN [22], [23], [24], [25]. Our study also showed that the classification accuracy does not improve radically with feature selection and extraction as most classifiers performed well with the raw data. Moreover, accuracy does not decrease with an increase in the number of classes (clusters, *K* = 5 to 11). This creates a question about our scheme, whether feature selection using phase portraits was necessary. The answer depends on the criteria considered for detection because we also found that the greedy algorithm such as DT classification accuracy was sensitive to feature selection, extraction, and the number of classes. Therefore, if the goal is to detect different IBM sub-phases with high accuracy and ignore other aspects such as processing time and flexibility of using different algorithms, then feature selection and extraction using phase portraits are not necessary. However, ignoring these criteria will increase the hardware cost as faster processors and built-in computation methods will be needed. Therefore, our detection scheme offers high accuracy, speed, and flexibility with other algorithms without loss of accuracy (robustness).

The other aspect of the classification accuracy of our scheme is parameter optimization. We have mentioned in our results that we kept parameters consistent while comparing the accuracy with and without feature selection. Hence, the DT model without features was not optimized subjectively to attain higher accuracy. The rationale was to reduce any analysis bias. However, the biased approach of optimizing the parameter for higher accuracy of the DT training model will not change the fact that there will still be a trade-off between accuracy and speed. Therefore, in our study, feature selection and extraction using phase portraits are of significant importance as it reduces the constraints related to processing speed without impact accuracy [26], especially for greedy deterministic algorithms like DT.

### 4.3. Limitations & Future work

There are information-based constraints with two-dimensional features 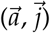 in this study. The lack of high-dimensional feature space limits the degree of freedom to process enough information. But additional information can be captured using multichannel sensors with a high sampling rate. The inertial measurement units (IMUs) sensors can be an alternate [17], [27]. The IMUs are embedded with accelerators, gyroscopes, and magnetometers, and they can provide linear and rotational kinematic information about IBM in the vertical direction [27], [28]. Therefore, a high-dimensional feature space can be acquired using such sensors and will be the scope of our future studies.

In addition, our study does not account for the relationship between physiological factors [29] (such as motivation level, blood lactic acid level, lung volume, partial levels of *O*_2_ and *CO*_2_ (*PO*_2_ and *PCO*_2_) and IBM sub-phases. Therefore, in this study, IBM’s physiological role is inferred. These physiological factors are the key factors for SP characteristics determination [7], [10]. Thus, we cannot develop a concrete relationship between IBM sub-phases and SP physiological characteristics. It is something we will be considering studying in the future.

### 5. Conclusion

We found in our study that IBMs during BH can be captured through force sensors. In this study, we developed a pattern recognition and classification scheme using estimated 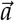 and 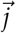 from the vertical ground reaction force. The 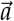 and 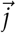 were used to quantify the rapid movements associated with involuntary contractions. Our scheme recognizes and classifies the IBM phase from the NM phase during BH, and also identifies and classifies different sub-phases of varying (low, moderate, and high) magnitude within the IBM phase. We also developed phase portraits using 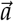 and 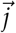 to select and extract specific features so that accuracy, robustness, and computational cost of our scheme can be enhanced. To the best of our knowledge, this is the first study that had developed a pattern recognition and classification scheme to detect IBMs from vertical ground reaction force.

## 6. Authors Declaration

### 6.1. Authors contribution

The study was conceived and designed by RES, GW, JMF, NPB, BA. JMF did the data collection. RES designed the scheme. RES and SG did the data analysis. RES drafted the manuscript. GW, NPB, JMF, and BA provided input and feedback on the study. All authors provided critical revision of the manuscript and final approval of the version to be published.

### 6.3. Ethical approval

The study was approved by the Colorado Mesa University, IRB # 22-35. The participants volunteered for the study with a signed consent, and all applicable institutional and governmental regulations concerning the ethical use of human participants were followed during the course of this study.

### 6.4. Data availability

The data is available from Dr. Gannon White and/or Dr. Nate P. Bachman on reasonable request. A simple R code for the recognition, classification using processed data with relevant packages is available below. Note:- The code is for processed data and uses test data to validate model accuracy. https://github.com/rajatsingh91/Classification-Recognition-Convulsion

## 6.5 Acknowledgment

We would like to thank Dr. Taimoor Afzal for his feedback on this manuscript and the research work. We also like to thank mdpi biomechanics for inviting us and providing waiver for this article.

